# A deep learning algorithm for 3D cell detection in whole mouse brain image datasets

**DOI:** 10.1101/2020.10.21.348771

**Authors:** Adam L. Tyson, Charly V. Rousseau, Christian J. Niedworok, Sepiedeh Keshavarzi, Chryssanthi Tsitoura, Lee Cossell, Molly Strom, Troy W. Margrie

## Abstract

Understanding the function of the nervous system necessitates mapping the spatial distributions of its constituent cells defined by function, anatomy or gene expression. Recently, developments in tissue preparation and microscopy allow cellular populations to be imaged throughout the entire rodent brain. However, mapping these neurons manually is prone to bias and is often impractically time consuming. Here we present an opensource algorithm for fully automated 3D detection of neuronal somata in mouse whole-brain microscopy images using standard desktop computer hardware. We demonstrate the applicability and power of our approach by mapping the brain-wide locations of large populations of cells labeled with cytoplasmic fluorescent proteins expressed via retrograde trans-synaptic viral infection.

## Introduction

To understand the circuits underlying computations in the brain, it is necessary to map cell types, connections and activity across the entire structure. Advances in labelling (1–3), tissue clearing (4–6) and imaging (7–12) now allow for the meso-and microscopic study of brain structure and function across the rodent brain. Analysis of these whole-brain images has lagged behind the developments in imaging (13). Although there are many relevant commercial and open-source bio-image analysis packages available (14–17), these have traditionally been developed for 2D images or for 3D volumes much smaller than a rodent brain.

In rodent studies, an increasingly common whole-brain image analysis task is the identification of individual, labelled cells across the entire brain. Traditionally, this was carried out manually (18–21), but this approach does not scale to all biological questions, particularly when many cells are labelled. Considering that a mouse brain has around 100 million neurons (22), even if only 0.01% of cells in the brain are labelled, a manual approach becomes impractical for any kind of routine analysis.

There are many methods that work well for identifying labelled cells in serial 2D sections, and subsequently registering the images to reference atlases (23–31). However, 2D analysis can be subject to bias as detected cell numbers can be under, or overestimated depending on sampling in the third dimension. There are now methods for 3D cell detection in whole-brain microscopy images (32–34), but these methods are limited to either nuclear labels, or were only validated in small regions of the brain. Although nuclear labels are much simpler to detect than membrane or cytoplasmic markers (as they have a simple shape and can be approximated as spheres and are far less likely to be overlapping in the image) there are many applications in which a nuclear label is not practical or even useful, as in the case of in vivo functional imaging. Any approach must also be validated throughout the brain, as the signal to noise (SNR) characteristics can vary between brain regions (e.g. sources of noise could be falsely detected as a cell). There does not yet exist a quick method for 3D detection of cells with cytosolic labels that has been validated throughout an entire brain. Meeting this need is a highly desired goal within systems neuroscience.

To overcome the limitations of traditional computer vision, machine learning — and particularly deep learning (35) — has revolutionised the analysis of biomedical and cellular imaging (36). Deep neural networks (DNNs) now represent the state of the art for the majority of image analysis, and have been applied to analyse whole-brain images, to detect cells in 2D (23, 28) or to segment axons (37). However, they have two main disadvantages when it comes to 3D whole brain analysis. Firstly, they require large amounts of manually-annotated training data (e.g. for cell segmentation, this would potentially require the painstaking annotation of hundreds or thousands of cell borders in 3D). Secondly, the complex architecture of DNNs means that for big data (e.g. whole-brain images at cellular resolution), large amounts of computing infrastructure is required to train these networks, and then process the images in a reasonable time frame.

To harness the power of deep learning for 3D identification of labelled cells in whole-brain images, we developed a computational pipeline which uses classical image analysis approaches to detect potentially labelled cells with high sensitivity (cell candidates), at the expense of detecting false positives (i.e. geometrically similar objects). This is then followed by application of a DNN to classify cell candidates as either true cells, or artefacts to be rejected. Harnessing the power of deep learning for object classification rather than cell segmentation at a voxel level speeds up analysis (since there are billions of voxels, but many fewer cell candidates) and simplifies the generation of training data. Rather than annotating cell borders in 3D, cell candidates from the initial step can be further classified by the addition of a single (cell or artefact) label.

## Results

To illustrate the problem and to demonstrate the software, whole mouse brain images were acquired following retrograde rabies virus labelling. Viral injections were performed into visual or retrosplenial cortex, causing thousands of cells to be cytoplasmically labelled throughout the brain. Data was acquired using serial two-photon microscopy as previously described (19) (Fig. 1). Briefly, coronal sections are imaged at high-resolution (2 µm × 2 µm × 5 µm voxel size) and stitched to provide a complete coronal section. This is carried out for ten imaging planes after which a microtome removes the most superficial 50 µm of tissue and the process is repeated until the entire brain data set is collected. Light emitted from the specimen is filtered and detected via at least two channels, a primary signal channel containing the fluorescence signal from labelled target neurons and a secondary ‘autofluorescence’ channel that does not contain target signals but provides anatomical outlines. An example single-plane image is shown in Fig. 2a.

**Fig. 1.**
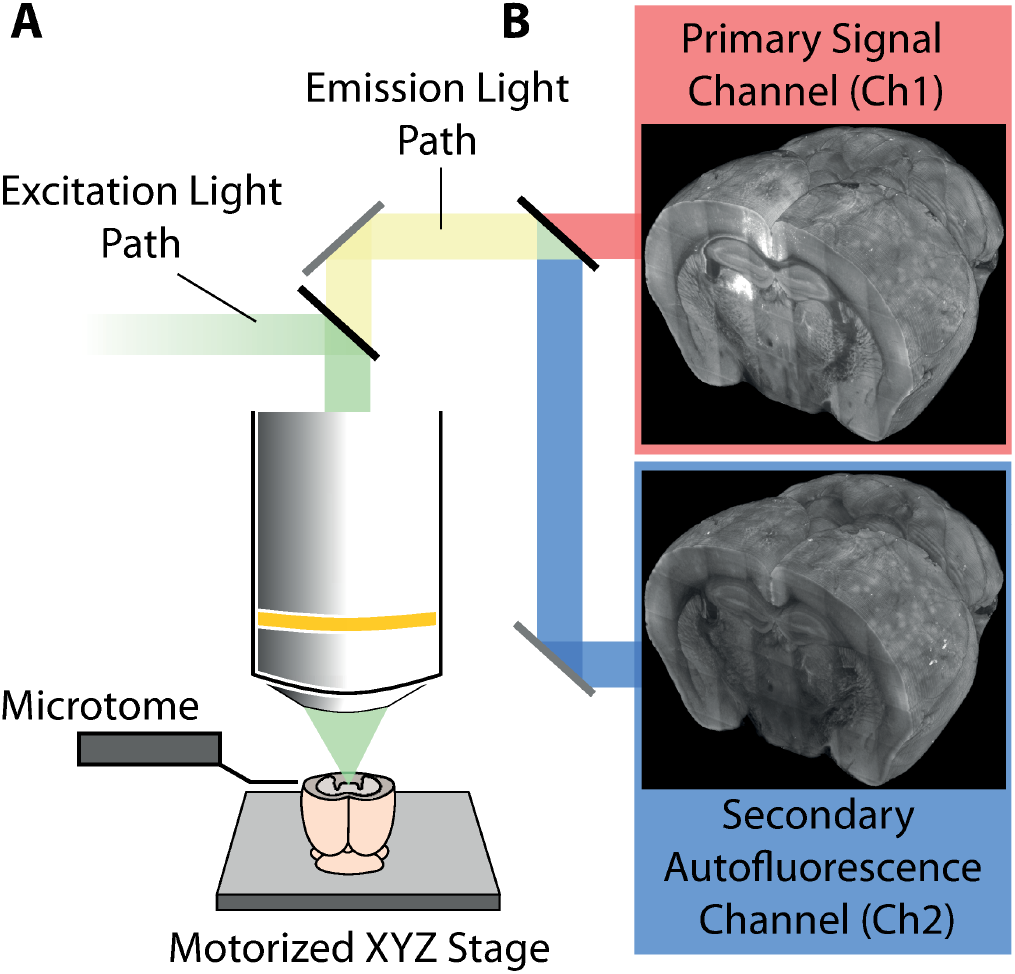
Simplified schematic diagram of the serial two-photon microscope and data acquisition process. A: The tissue is excited using a femtosecond Ti-sapphire laser (emission wavelength = 800 nm). For data collection, 50 µm of tissue (at approximately 40 µm to 90 µm below the tissue surface) is imaged in ten, 5 µm thick planes. An in-built microtome then physically removes a 50 µm thick section from the optical face. This process is repeated to generate a complete 3D dataset of the specimen. B: For signal collection, the emitted lightpath is split into two channels whereby the primary channel detects the fluorescence signal of interest from labelled cells (e.g. mCherry at 610 nm) and the second channel (e.g. at 450 nm) detects the tissue autofluorescence signal that reveals gross anatomical structure.

**Fig. 2.**
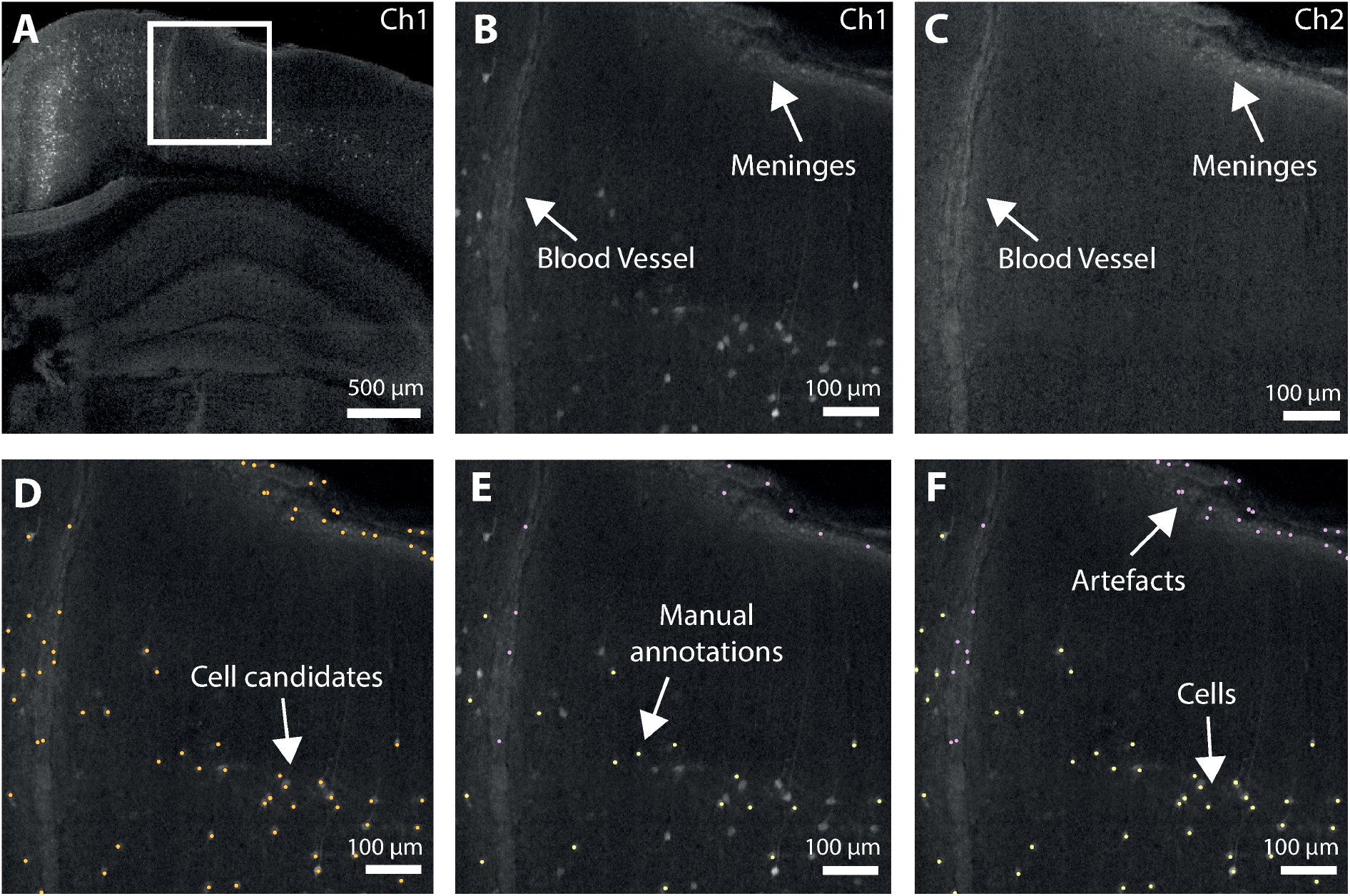
Illustration of the cell detection process. A: Single coronal plane of raw data (primary signal channel, Ch1). B: Enlarged insert of cortical region from A showing examples of structural features (artefacts) often erroneously detected. C: Cortical region shown in B, in the secondary autofluorescence channel (Ch2). Cells can only be seen in Ch1, but artefacts are visible in both channels. D: Detected cell candidates overlaid on raw data. Labelled cells as well as numerous artefacts are detected. E: Illustration of training data. A subset of detected cell candidates are classified as cells (yellow) or artefacts (purple). Cuboids of data centered on these selected cell candidates are then used to train the network. F: Classified cell candidates. The trained cell classification network is applied to all the cell candidates from (E) and correctly rejects the initial false positives.

### Cell candidate detection

When developing any object detection algorithm, a balance must be struck between false positives and false negatives. In traditional (two-dimensional) histology, simple thresholding (e.g. (38)) can often work well for cell detection. This does not necessarily apply to whole brain images. In samples with bright, non-cellular structures (artefacts, Fig. 2c) or lower signal to noise ratio, simple thresholding can detect many non-cellular elements. Image preprocessing and subsequent curation of detected objects can overcome some of these issues, but no single method works reliably across the brain in multiple samples. Either some cells are missed (false negatives), or many artefacts are also detected (false positives). To overcome this, a traditional image analysis approach was used to detect cell candidates, i.e. objects of approximately the correct brightness and size to be a cell. This list of candidates is then later refined by the deep learning step. Crucially, this refinement allows the traditional analysis to produce many false positives while minimising the number of false negatives. Images are median filtered and a Laplacian of Gaussian filter is used to enhance cell-like objects. The resulting image is thresholded, and objects of approximately the correct size are saved as candidate cell positions (Fig. 2d). An overview of the cell candidate detection steps is shown in Fig. S1. The thresholding is tuned to pick up every detectable cell but this also results in the detection of many false positives that often appear as debris on the surface of the brain and in some cases unidentified objects within blood vessels. This initial detection step is based on a number of tunable parameters (Table 5) which were chosen based on trial and error to be biased to over detection, and so be relatively robust to changes in the input data. They may of course need to be tuned for very different data, in particular the in-plane cell somata diameter which may need to be tuned when different types of cells are labelled.

**Table 5.** Default cell detection parameters.

| Parameter | Value |
| --- | --- |
| In-plane cell somata diameter (∅) | 16 $\mu\text{m}$ |
| Gaussian smoothing sigma | 0.2∅ |
| Intensity threshold | Mean + 10 x SD |
| In-plane (lateral) ellipsoidal filter width | 6 $\mu\text{m}$ |
| Axial (z) ellipsoidal filter width | 15 $\mu\text{m}$ |
| Ellipsoidal filter overlap threshold fraction | 0.6 |

### Cell candidate classification using deep learning

A classification step, which uses a 3D adaptation of the ResNet (39) convolutional neural network (Figs. S2 & S3) is then used to separate true from false positives. To classify cell candidates, a subset of cell candidate positions were manually annotated (e.g. Fig. 2e). In total, ∼100,000 cell candidates (50,653 cells and 56,902 non-cells) were labelled from five brains. Small cuboids of 50 × 50 × 100 µm around each candidate were extracted from the primary signal channel along with the corresponding cuboid in the secondary autofluorescence channel (Fig. 3a). This allows the network to “learn” the difference between neuron-based signals (only present in the primary signal channel), and other non-neuronal sources of fluorescence (potentially present in both channels).

**Fig. 3.**
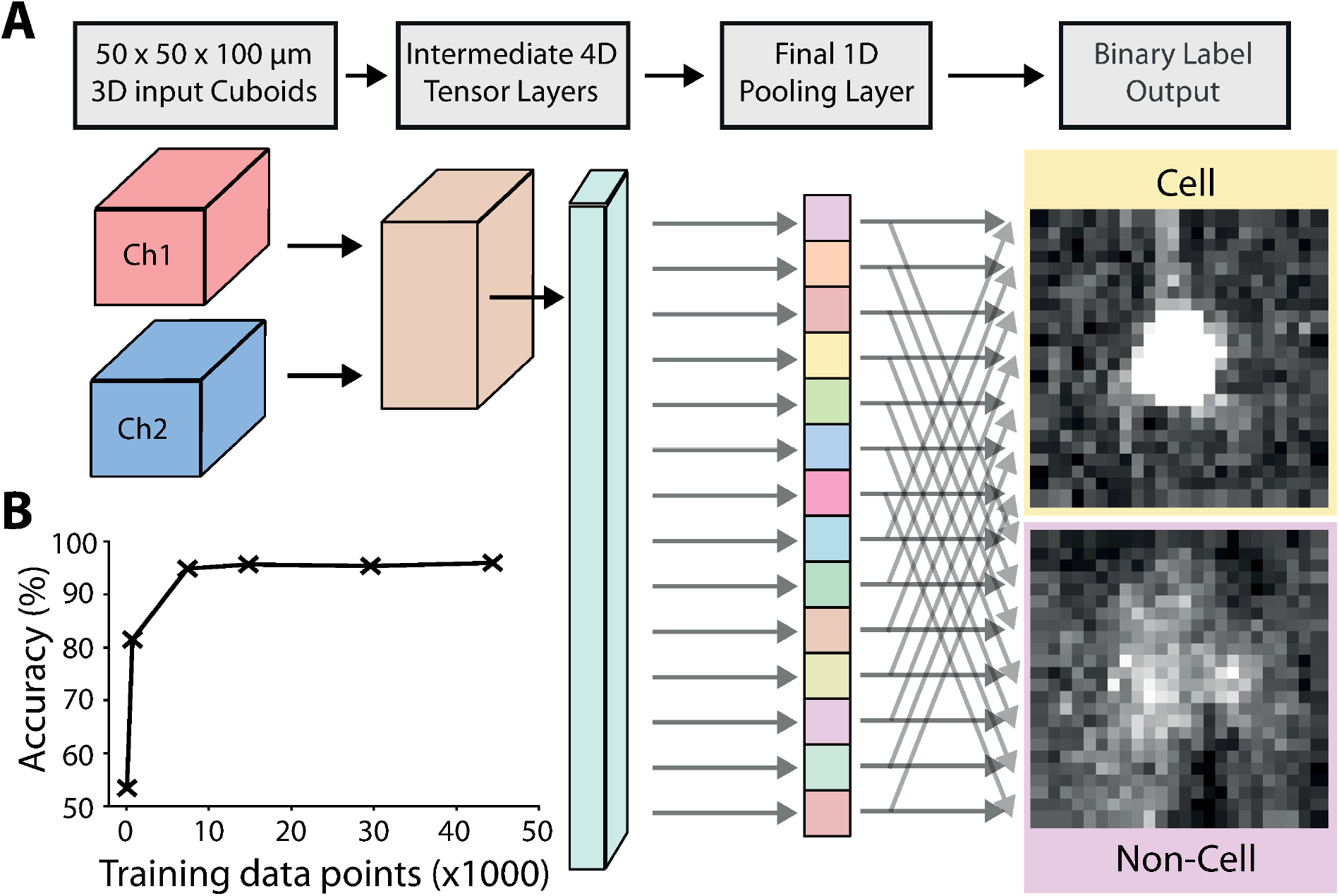
Cell classification. A: The input data to the modified ResNet are 3D image cuboids (50 µm × 50 µm × 100 µm) centered on each cell candidate. There are two cuboids, one from the primary signal channel, and one from the secondary autofluorescence channel. The data is then fed through the network, resulting in a binary label of cell or non-cell. During training the network “learns” to distinguish true cells, from other bright non-cellular objects. See Figs. S2 & S3 for more details of the 3D ResNet architecture. B: Training the initial cell classification network: classification accuracy as a function of training data quantity.

The trained classification network is then applied to classify the cell candidates from the initial detection step (Fig. 2f). The artefacts (such as those at the surface of the brain and in vessels) have been correctly rejected, while correctly classifying the labelled cells. To quantify the performance of the classification network, and to assess how much training data is required for good performance, the manually annotated training data was split up into a new training dataset from four brains, and a test dataset from the fifth brain. A new network was trained on subsets of the training data, and performance tested on the fifth brain (15,872 cells and 18,168 non-cells). Fig. 3b shows that relatively little training data was required for good performance on unseen test data, with 95% of cell candidates classified correctly with ∼7,000 annotated cell candidates. Although ∼7,000 data points are required to train the network from scratch, we provide the network trained on the full dataset with the software. Users can then re-train this network with a much smaller amount of experiment-specific training data.

### Application

To illustrate the method, the cell detection software was applied to data which was not used to develop or train the classification network. Data from a previous experiment (40) was acquired on a different microscope and was used to simulate real-world usage in which the SNR characteristics of the data may vary from that used to pre-train the supplied network. Neurons presynaptic to layer 2/3 primary visual cortical cells were labeled using rabies virus tracing (expressing mCherry) in two Penk-Cre mice.

The algorithm was run on these two brains, using the default candidate detection parameters (Table 5). A small number (619) of the detected cell candidates were manually annotated on a single brain (brain 1) including confirmations of correct classifications, and corrections of incorrect classifications. The pre-trained network was then retrained using these data points and 10% of the data was held back for validation during training. The network was trained until the validation loss function began to plateau, taking 73 minutes. The full cell detection algorithm was then repeated for both brains using the re-trained network on a laptop computer. Total time for cell detection was 83 minutes for brain 1 and 91 minutes for brain 2 (for full timings see Table 1).

**Table 1.**
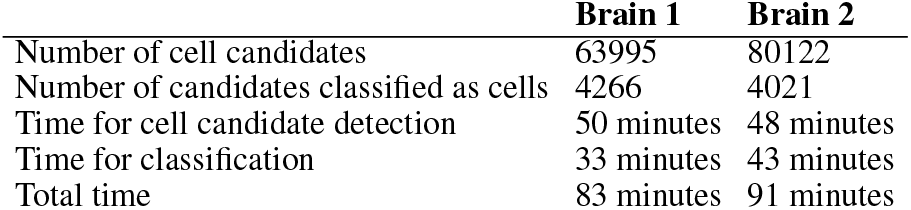
Algorithm timings on a laptop computer with Intel i9-9900K CPU, 32GB RAM and an NVIDIA RTX2080 GPU. Data stored on an external solid-state drive.

|  | Brain 1 | Brain 2 |
| --- | --- | --- |
| Number of cell candidates | 63995 | 80122 |
| Number of candidates classified as cells | 4266 | 4021 |
| Time for cell candidate detection | 50 minutes | 48 minutes |
| Time for classification | 33 minutes | 43 minutes |
| Total time | 83 minutes | 91 minutes |

To assign detected cells to a brain region, the Allen Mouse Brain Reference Atlas (ARA (41)) annotations were registered to the secondary autofluorescence channel using brainreg (42), a Python port of the validated aMAP pipeline (43). These annotations were overlaid on the raw data (Fig. 4a), and the number of cells in each brain region were reported, allowing for quantitative analysis (Table 2).

**Fig. 4.**
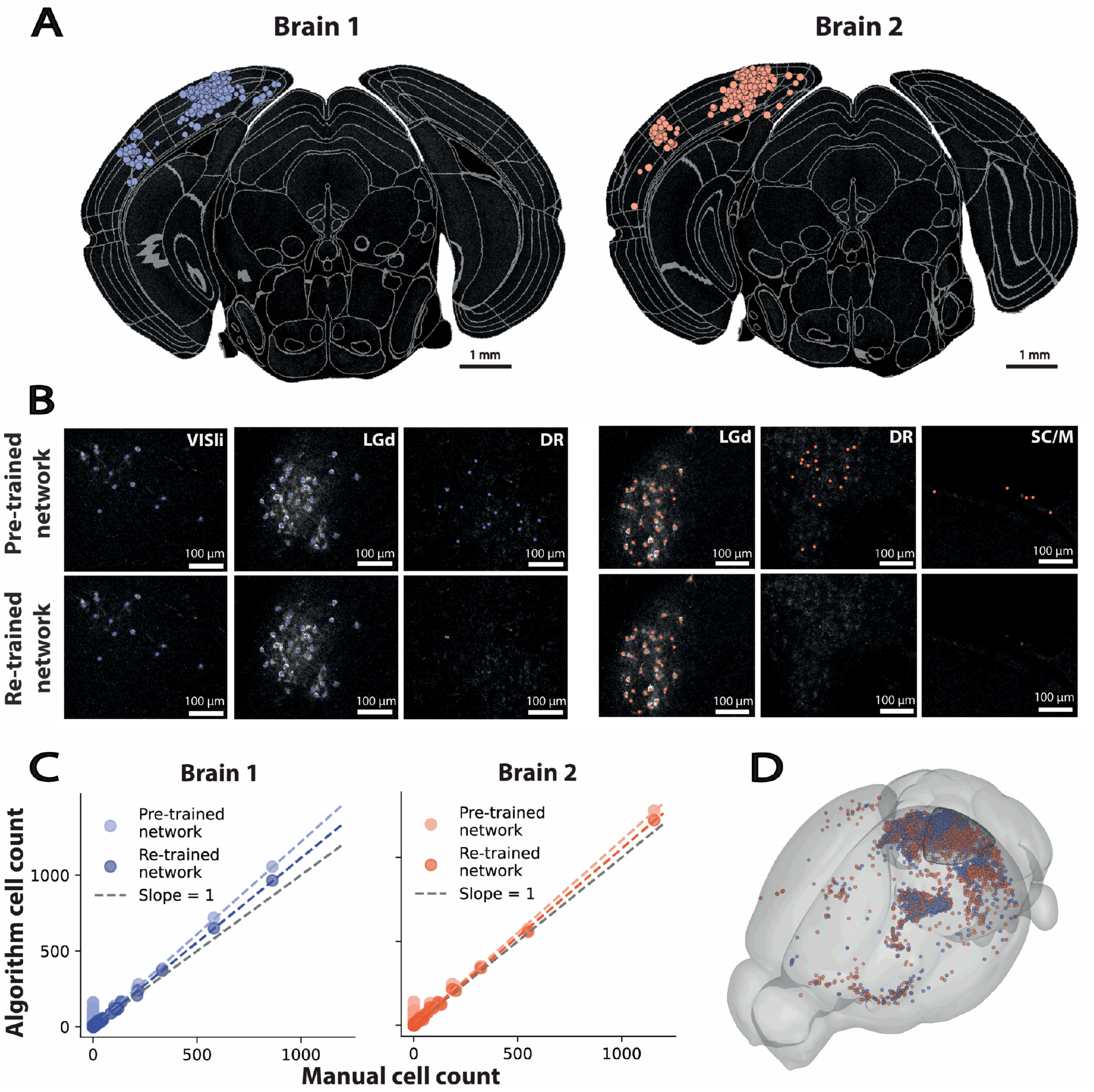
Application of the algorithm to unseen data. Presynaptic neurons labelled by rabies viral injection into layer 2/3 primary visual cortex in Penk-Cre mice. A: Detected cells overlaid on raw data along with the brain region segmentation for Brain 1 (blue) and 2 (orange). The size of the coloured disk represents the proximity of the cell centroid to the image plane displayed. B: Comparison of cell detection before and after re-training the pre-trained network in different regions of the image. Cells with different morphologies are correctly detected in both dense and sparse regions, and artefacts are rejected. VISli — Laterointermediate area, LGd — Dorsal part of the lateral geniculate complex, DR — Dorsal nucleus raphe, SC/M — Superior colliculus & meninges. C: Comparison of cell counts per ARA brain region between the algorithm and the mean of the two expert counts. Best fit shown before and after re-training. D: Visualisation of detected cells from both brains warped to the ARA coordinate space in 3D, along with the rabies virus injection site target (primary visual cortex, wireframe).

**Table 2.** Total number of cells in each region, per brain, projecting to layer 2/3 neurons in primary visual cortex. Ten regions with the greatest number of cells across both brains shown.

| Brain structure name | Brain 1 | Brain 2 |
| --- | --- | --- |
| Primary visual area, layer 2/3 | 965 | 1222 |
| Primary visual area, layer 5 | 650 | 558 |
| Dorsal part of the lateral geniculate complex, core | 371 | 340 |
| Lateral posterior nucleus of the thalamus | 240 | 215 |
| Primary visual area, layer 4 | 207 | 205 |
| Retrosplenial area, ventral part, layer 5 | 162 | 135 |
| Dorsal part of the lateral geniculate complex, shell | 122 | 124 |
| Lateral dorsal nucleus of thalamus | 122 | 107 |
| Retrosplenial area, dorsal part, layer 5 | 110 | 103 |
| Posteromedial visual area, layer 5 | 46 | 74 |

The new data was noisier than that used to pre-train the network (possibly due to the use of resonant vs galvanometer scanning), which lead to false-positives throughout the brain. A small amount of new training data, taking approximately five minutes to generate was sufficient to significantly improve performance of the algorithm (Fig. 4b). The re-trained network removed many of the false-positives, while still correctly classifying the labelled cells.

### Validation

To quantify the accuracy of the algorithm brain-wide, we generated ground truth data for both the brain used to generate data for re-training (brain 1) and the “unseen” brain (brain 2). Two experts manually annotated all labelled cell somata throughout the brains. These cells were assigned to regions in the ARA in the same way as the automated cell counts and an average of the two experts was taken. The comparison between the automated counts and the manual counts is shown in Fig. 4c. Using the pre-trained network, the algorithm detects false positives, including in many areas with no labelled cells. Re-training the network significantly reduces the number of false positives, bringing the best-fit line closer to an exact match to the manual cell counts. For both brains, a linear fit to the algorithm and manual cell counts is above 1 (brain 1 - 1.113, brain 2 - 1.053), suggesting a small number of false positives (Table 3).

**Table 3.** Comparison between algorithm and expert cell counting.

|  | Brain 1 |  |
| --- | --- | --- |
|  | Pearson correlation coefficient | Linear best-fit slope |
| Algorithm vs expert 1 | 0.999 | 1.054 |
| Algorithm vs expert 2 | 0.999 | 1.178 |
| Expert 1 vs expert 2 | 0.999 | 1.116 |
| Algorithm vs expert mean | 0.999 | 1.113 |
|  | Brain 2 |  |
|  | Pearson correlation coefficient | Linear best-fit slope |
| Algorithm vs expert 1 | 0.998 | 0.964 |
| Algorithm vs expert 2 | 0.997 | 1.151 |
| Expert 1 vs expert 2 | 0.994 | 1.188 |
| Algorithm vs expert mean | 0.999 | 1.053 |

Although the results from the two brains look similar when the detected cells are warped to the ARA coordinate space (Fig. 4d), there is still significant biological variability. For this reason, most experiments quantify relative, rather than absolute, cell counts (18, 20, 24, 44). It is therefore important than the correlation between the automated cell counts and ground truth is as high as possible. The correlation for both the brain used for training (brain 1), and the “unseen” brain (brain 2) is very high (Pearson correlation coefficient, *ρ* = 0.999), and higher than the correlation between the automated cell counts for both brains (*ρ* = 0.982).

### Effect of varying axial sampling

All the data presented was acquired with high axial sampling (5 µm), but this is not always possible or desirable due to imaging time, data storage requirements, or biological considerations such as photobleaching. To assess the performance of the algorithm with varying optical slice thicknesses, we downsampled the data from brain 2, to generate synthetic datasets with axial sampling of 5, 10, 20 and 40 µm, and the algorithm was applied as before.

The cell counts were compared to the mean of expert counts (Table 4). Performance in terms of absolute cell numbers (best fit line slope, and Pearson correlation coefficient) was comparable for 5, 10 and 20 µm, although the 5 µm dataset was most highly correlated to the ground truth. The performance on the 40 µm dataset was much worse. The best-fit line slope of 0.675 means that many cells have been missed, although the high correlation (*ρ* = 0.942) suggests that this effect is relatively uniform throughout the brain.

**Table 4.** Algorithm performance compared to mean of expert manual counts with varying axial sampling.

| Axial sampling [$\mu\text{m}$] | Pearson correlation coefficient | Linear best-fit slope |
| --- | --- | --- |
| 5 | 0.999 | 1.053 |
| 10 | 0.978 | 0.906 |
| 20 | 0.981 | 1.011 |
| 40 | 0.942 | 0.675 |

The results are not surprising, as even with 20 µm axial sampling, most cells can still be visualised in multiple image planes, and so 3D cell detection is still possible. At 40 µm axial sampling, this is not possible, and so many cells are missed. Although the correlation coefficients at 10 and 20 µm axial spacing are relatively high (*ρ* = 0.978, 0.981), they are lower than the correlation between different brains (*ρ* = 0.982). This may reduce the likelihood of detecting small biological effects using data with low axial sampling. There are also many other factors that affect how many image planes a cell will appear in, such as the cell somata size, and the axial point spread function of the microscope.

## Discussion

Mapping the distribution of labelled neurons across the brain is critical for a complete understanding of the information pathways that underlie brain function. Many existing methods for cell detection in whole-brain images rely on classical image processing, which can be affected by noise, and may not detect complex cell morphologies. DNNs can be used for highly sensitive image processing, but often require laborious generation of training data and are prohibitively slow for the analysis of large, 3D images. The presented method here overcomes these limitations by combining traditional image processing methods for speed, with a DNN to improve accuracy.

Recent developments in microscopy technology (e.g. (12)) now allow for quicker, more routine acquisition of whole-brain datasets. It is important that the image analysis can be carried out in a timely fashion, and without relying on large-scale computing infrastructure. Processing time for the ∼180GB images in Fig. 4 on a laptop was around 90 minutes, so sixteen datasets could be analysed in a single day, much quicker than the sample preparation and imaging steps. Once parameters are optimised, and the classification network is trained, the software can run entirely without user intervention.

In traditional DNN approaches for image analysis, generation of training data is often a major bottleneck. While large-scale “citizen science” approaches can be used to generate large amounts of training data (45), this is not practical for the majority of applications, e.g. when anatomical expertise is required. Our method overcomes this by requiring only a binary label (cell or non-cell) for each cell candidate in the training dataset, rather than a painstaking 3D outline of each cell. The software is released with a pre-trained network, and so the network can be re-trained for specific datasets very quickly. The total time to generate the new network used to classify images in Fig. 4 was less than two hours, including generating training data and retraining the network.

We show that the results of the proposed method compare well to expert manual counts, particularly the correlation between counts in different brain areas. However our results also show the importance of re-training the pre-trained network for new datasets (even if they are superficially similar). It is important to also bear in mind that although we quantified performance of the algorithm across the brain, we did not label every type of cell, and some areas of the brain with densely packed neurons (e.g. hippocampus) were only sparsely labelled. When applying this method to very different data, users should ensure that they re-train the model with representative training data (including different brain areas, cell types and image artefacts if applicable), and check the results in detail. The data in Table 4 shows that although the method is relatively robust to the axial sampling, true 3D imaging (i.e. labelled cells appear in at least two axial planes) is required for accurate cell detection using our method.

The ability to quickly detect, visualise and analyse cytoplasmically labelled cells across the mouse brain brings a number of advantages over existing methods. Analysing an entire brain rather than 2D sections has the potential to detect many more cells, increasing the statistical power and the likelihood of finding novel results, particularly when studying rare cell types. Whole-brain analysis may also be less biased than analysing a series of 2D planes, especially in regions with low cell densities, or differing cell sizes.

This software is fully open-source, and has been written with further development and collaboration in mind. In future we aim to adapt the network to be flexible as to the number of input channels, and output labels. The classification network currently relies on using both the primary signal and the secondary autofluroescence channel. In the future it would be valuable to train a network that could achieve a similar level of performance using a single input channel. Analysing a single channel would allow half as much data to be collected (although autofluorescence channels are optimal for atlas registration). Training a network to produce multiple labels (rather than just cell or non-cell) would allow for cell-type classification based on morphology, or based on gene or protein expression levels if additional signal channels were supplied. Although the ResNet architecture was chosen based on performance (39) and flexibility in new contexts (e.g. (46)), there are many newer network architectures that could be implemented to improve performance (e.g. (47, 48)). Lastly, although this approach was designed for fast analysis of large whole-brain datasets, the proposed two-step approach could be used for any kind of large-scale 3D object detection.

## DATA AVAILABILITY

The methods outlined in this manuscript are available within the cellfinder software, part of the BrainGlobe suite of computational neuroanatomy tools. The software is open-source, written in Python 3 and runs on standard desktop computing hardware (although a CUDA compatible GPU allows for a considerable reduction in processing time). Source code is available at github.com/brainglobe/cellfinder and pre-built wheels at pypi.org/project/cellfinder. Documentation, tutorials, and the data underlying Fig. 4 are available at docs.brainglobe.info/cellfinder.

## ACKNOWLEDGEMENTS

This work was supported by grants from the Gatsby Charitable Foundation (GAT3361) and Wellcome Trust (090843/F/09/Z and 214333/Z/18/Z) to T.W.M..This manuscript was typset using a modified version of the HenriquesLab bioRxiv template.

## Materials and methods

All experiments were carried out in accordance with the UK Home Office regulations (Animal Welfare Act 2006) and approved by the establishments Animal Welfare and Ethical Review Board.

### Sample preparation

All mice used were transgenic Cre-reporter (Ntsr1-Cre, GAD2-IRES-Cre, Rbp4-Cre & PenkCre) mice bred on a C57BL/6 background. The mice were anesthetized and an AAV Cre-dependent helper virus encoding both the envelope protein receptor and the rabies virus glycoprotein was stereotactically injected into visual cortex or retrosplenial cortex. Four days later, a glycoprotein deficient form of the rabies virus expressing mCherry was delivered into the same site. After ten further days, the animal was deeply anaesthetized and transcardially perfused with cold phosphate buffer (0.1 M) followed by 4% paraformaldehyde (PFA) in PB (0.1 M) and the brain left overnight in 4% PFA at 4 °C.

### Imaging

All data was acquired using serial section two-photon tomography (8). To generate the data to pre-train the deep-learning model, fixed brains were embedded in 4% agar and placed under a two-photon microscope containing an integrated vibrating microtome and a motorized x-y-z stage as previously described (19). Coronal images were acquired via two optical pathways (red and blue) as a set of 6 by 9 tiles with a voxel size of 1 µm × 1 µm obtained every 5 µm using an Olympus 10x objective (NA = 0.6) mounted on a piezo-electric element (Physik Instrumente, Germany). Following acquisition, image tiles were corrected for uneven illumination by subtraction of an average image from each physical section. Tiles were then stitched using a custom FIJI (14) plugin (modified from (49)) and downsampled to 2 µm × 2 µm × 5 µm voxel size.

To generate the data to test the algorithm, data was acquired using a different, custom-built resonant-scanning system controlled by ScanImage (v5.6, Vidrio Technologies, USA) using BakingTray (50), a custom software wrapper for setting up the imaging parameters. Images were assembled using StitchIt (51). Both brains were imaged in a single acquisition using a Nikon 16x objective (NA = 0.8), with a voxel size of 2.31 µm × 2.31 µm × 5 µm.

### Cell candidate detection

To detect cell candidates (broadly defined as anything of sufficient brightness and of approximately the correct size to be a cell), initially data from the primary signal channel was processed in 2D (Fig. S1). Images were median filtered, and then a Laplacian of Gaussian filter was performed to enhance small, bright structures (e.g. cells). This filtered image was binarised using a threshold calculated for each image plane (mean of image plane + 10 × image plane standard deviation). The thresholded image was then passed to an ellipsoidal filter to remove noise. Every position of this spatial filter in which the majority (given by an input parameter, ellipsoidal filter overlap threshold fraction) of the filter overlaps with thresholded voxels was saved as a potential cell candidate. This is used to remove noise (from e.g. neurites).

All cell candidates that form continuous spatial structures were merged together, and classed as a single cell candidate. If the resulting cluster was too large to be a single cell (based on the input cell somata size parameter), then this cluster was split into individual cell candidates using an iterative ellipsoidal filter. Briefly, the ellipsoidal filter was applied to all voxels within the cell candidate cluster, and any resulting cell candidate coordinate positions were recorded. The thresholded image was eroded, and the filter was reapplied on the eroded set of candidate voxels. This process was repeated for ten iterations, or until there were no cell candidates remaining. This process ensures that densely labelled cells are split into individual cell candidates.

Once the final list of candidate cells is determined, the centroid of each cell candidate (based on the 3D mean coordinate of thresholded voxels) was calculated, and the coordinates were saved as an XML file. All of these steps were all carried out using the default software parameters (Table 5), with the exception of the 40 µm axial spacing dataset for which the axial extent of the ellipsoid filter was increased to 30 µm.

### Cell candidate classification using deep learning

Cell candidates were classified using a ResNet (39), implemented in Keras (52) for TensorFlow (53). 3D adaptations of all networks from the original paper are implemented in the software (i.e. 18, 34, 50, 101 and 152-layer) and can be chosen by the user, but the 50-layer network was used throughout this study. The general architecture of these networks is shown in Figs. S2 & S3.

To generate data to train the classification network, output from the candidate detection step (cell candidate coordinates) were manually classified using a custom FIJI (14) plugin, or an integrated tool (cellfinder-curate) using napari (54) that is supplied with the software. Expert annotators were presented with the raw data, with cell candidates represented by hollow spheres, centered on the cell candidate. By scrolling through the 2D planes of the 3D dataset, the experts could view the cell candidate in 3D before marking it as a cell or artefact. Candidates were determined to be cells or artefacts based on size, shape (including the presence of neurites) and the fluorescence level compared to the secondary autofluorescence channel (which was visualised at the same time). Candidates were labelled by three experts, labelling different brains, and a subset of labelled candidates were cross-checked between experts.

Image cuboids of 50 µm × 50 µm × 100 µm (resampled to 50 × 50 × 20 voxels) were extracted from both the primary signal, and secondary autofluorescence channels, centered on the coordinates of the manually classified cell candidate positions. To increase the size of the training set, data were randomly augmented. Each of the following transformations were applied with a 10% likelihood: (i) flipping in any of the three axes, (ii) rotation around any of the three axes (between 45° to 45°) and (iii) circular translation along any of the three axes (up to 5% of the axis length). The networks were trained using an NVIDIA TITAN RTX GPU with a batch size of 32 and the Adam (55) method was used to minimise the loss (categorical cross entropy), with a learning rate of 0.0001. Cell candidates were classified using the trained network, and saved as an XML file with a cell or artefact label.

### Image registration and segmentation

To allow detected cells to be assigned an anatomical label, and for them to be analysed in a common coordinate framework, a reference atlas (Allen Mouse Brain Atlas, ARA, (41), provided by the BrainGlobe Atlas API (56)) was registered to the autofluorescence channel. This was carried out using brainreg (42), a Python port of the automatic mouse atlas propagation (aMAP) software (43), which itself relies on the medical image registration library, niftyreg (57). Firstly the sample brain was downsampled to the same voxel spacing as the atlas (10 µm isotropic) and was reoriented to the atlas template brain. These two images were then filtered to remove-high frequency noise (greyscale opening and flat-field correction). The images were firstly registered using an affine transform (*reg_aladin* (58)), followed by freeform non-linear registration (*reg_f3d* (57)). The resulting transformation was applied to the atlas brain region annotations (and a custom hemispheres atlas) to bring it into the same coordinate space as the sample brain.

### Validation

To compare results of the algorithm to ground truth, two experts manually annotated each cell in two whole-brain images using the same cellfinder-curate tool used to generate the training data. The experts were shown the full-resolution images, with both channels displayed in different colors. The experts could scroll through the 3D images plane by plane, zoom in and out, and adjust contrast settings to best visualise cells in different brain areas. Experts annotated cells based on the same criteria as for generating the training data (shape, size and fluorescence intensity). Annotating a cell position displayed a hollow sphere on top of the image, around the cell coordinate. This was visible in multiple image planes to ensure that individual cells were not labelled more than once.

The cell classification network was retrained for the new datasets, and the algorithm was run with the default cell candidate detection parameters. The images were also registered to the ARA, and cell coordinates were assigned to brain regions for both the automated and manual cell counts. The different cell counting approaches were firstly compared by calculating the Pearson correlation coefficient using Pandas (59, 60). To assess the bias of the different approaches, they were compared by calculating the slope of the best fit line by fitting a linear model using scikit-learn (61).

### Effect of varying axial sampling

To generate synthetic datasets with varying axial sampling, a single brain was downsampled in 3D by selecting every Nth image plane. For example, to generate a dataset with 20 µm sampling from the original dataset sampled at 5 µm, every fourth plane was used.

### Visualisation

For visualisation of data in standard space, detected cells must be transformed to the atlas coordinate space. Firstly, the affine transform from the initial registration was inverted (using *reg_transform*). The sample brain was then registered non-linearly to the atlas (again using *reg_f3d*) and a deformation field (mapping points in the sample brain to the atlas) was generated (using *reg_transform*). This deformation field was applied to the coordinates of the detected cells for each sample, transforming them into atlas coordinate space.

Plots were generated using Matplotlib (62), and image visualisation was performed using napari (54) and brainrender (63).

## Supplementary figures

**Fig. S1.**
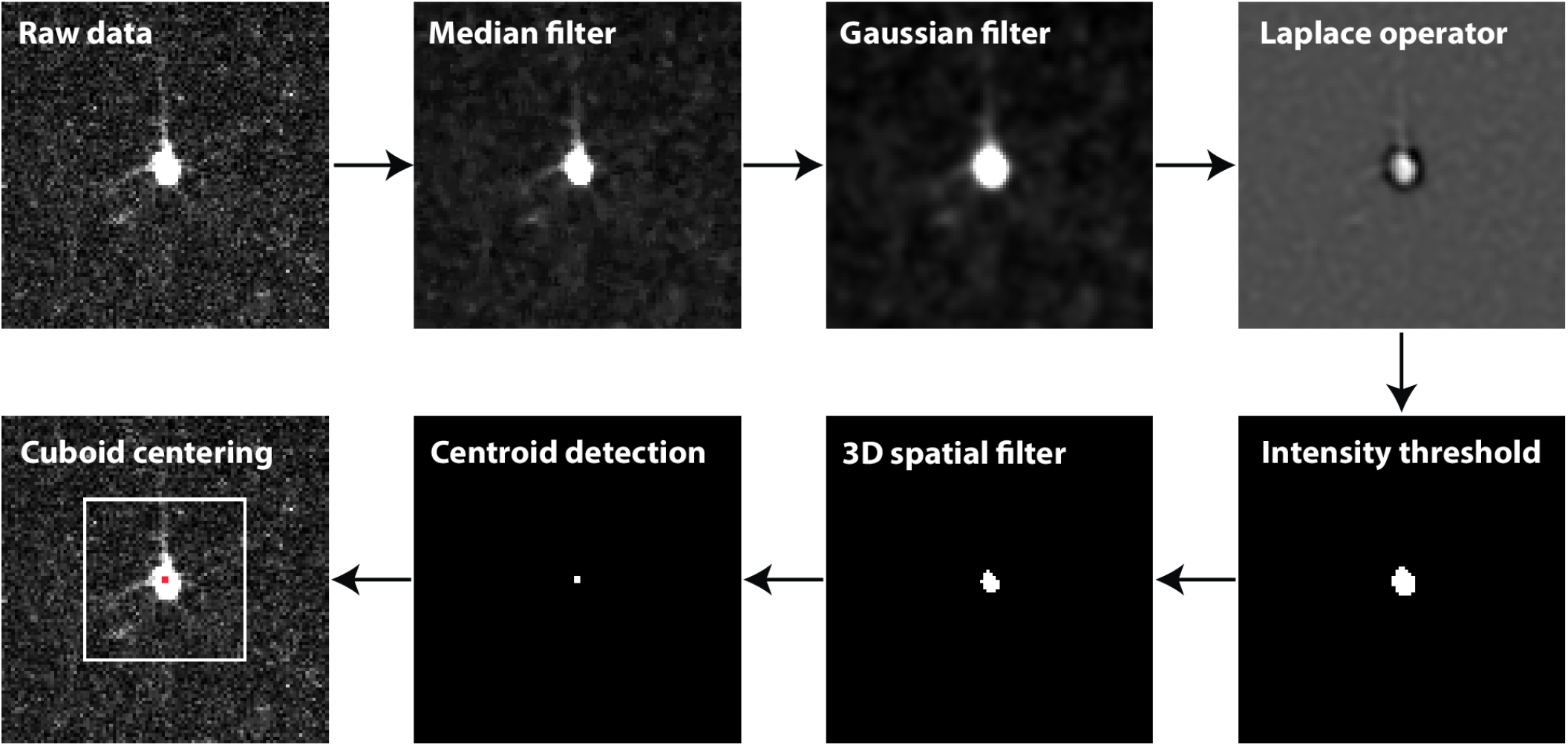
Overview of the initial cell candidate detection steps, from raw data to a cuboid of data fed into the classification network. Upper row: from left to right, the raw image is median filtered to remove noise. A Laplacian of Gaussian is then performed to enhance small, bright structures such as the cell soma. Lower row: from right to left, the image is thresholded and a 3D ellipsoidal filter is used to remove small, non-cellular objects (not shown in this image plane). The centroid of the resulting object is then used to center the cuboid of data that it passed to the deep learning classification network. Images shown are 100 µm × 100 µm, and the cuboid is 50 µm × 50 µm (and 100 µm in the third dimension).

**Fig. S2.**
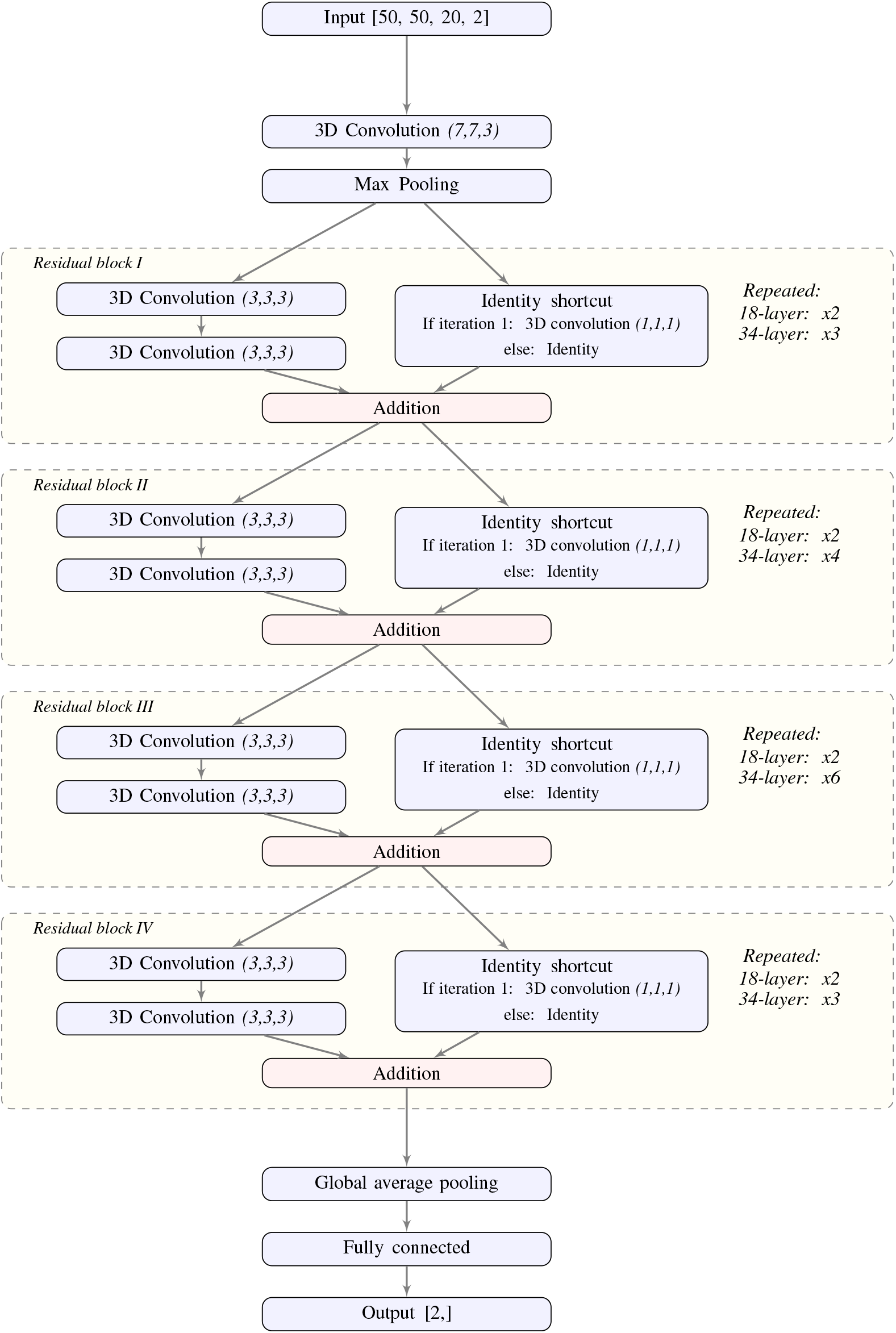
Architecture of the 3D ResNet. 3D adaptation of the 2D networks from [38] which are available for use in the software.

**Fig. S3.**
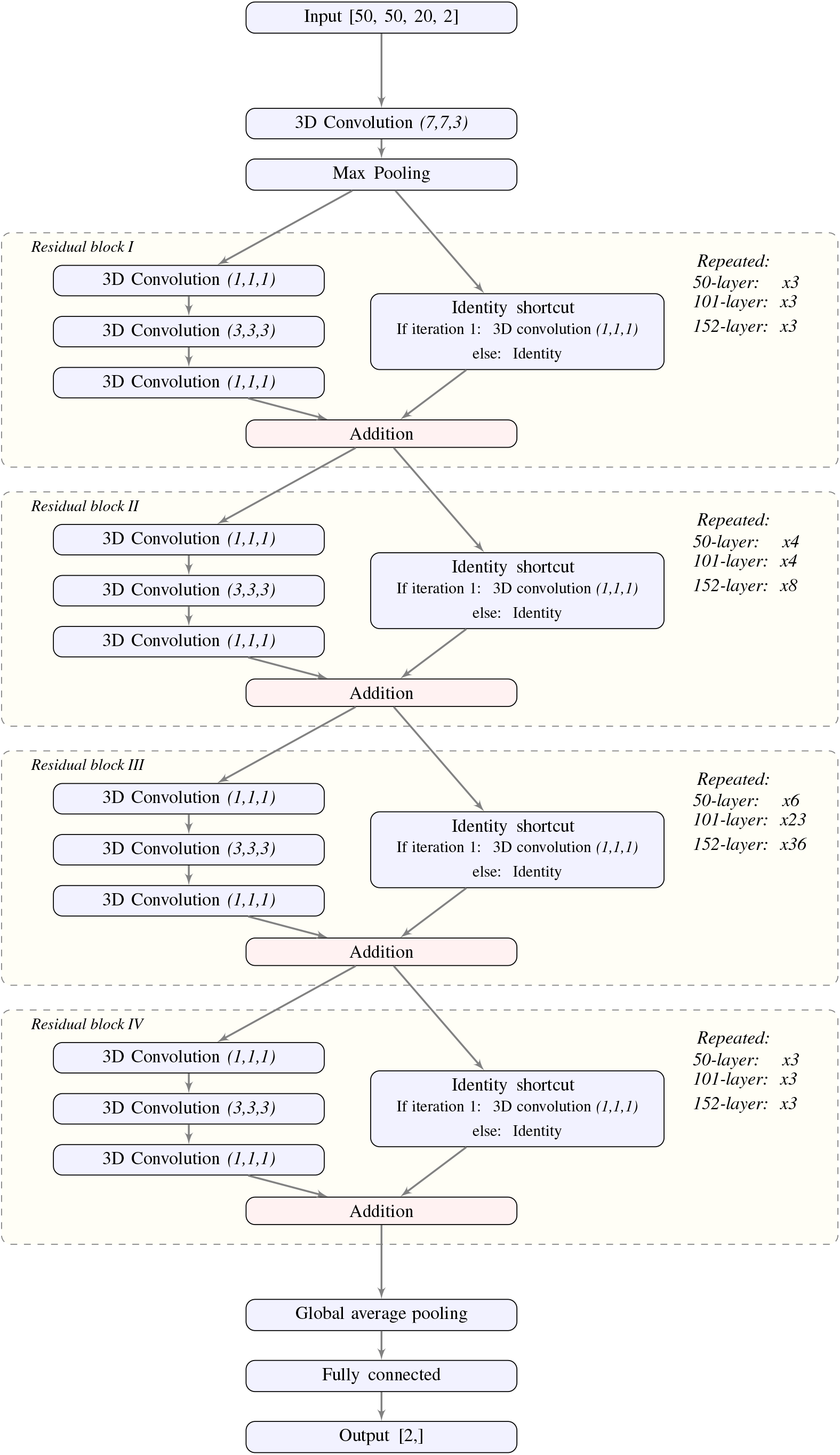
Architecture of the bottleneck 3D ResNet. 3D adaptation of the bottleneck 2D networks from [38] which are available for use in the software. The 50-layer bottleneck network is used throughout this study, and is used for the pre-trained model supplied with the software.

